# Selective attention controls olfaction in rodents

**DOI:** 10.1101/236331

**Authors:** Kaitlin S. Carlson, Marie A. Gadziola, Emma S. Dauster, Daniel W. Wesson

**Affiliations:** Department of Pharmacology & Therapeutics, University of Florida 1200 Newell Dr. Gainesville, FL, 32610. U.S.A; Center for Smell and Taste University of Florida 1200 Newell Dr. Gainesville, FL, 32610. U.S.A; Department of Neurosciences Case Western Reserve University 2109 Adelbert Rd. Cleveland, OH, 44106. U.S.A

## Abstract

Critical animal behaviors, especially among rodents, are guided by odors in remarkably well-coordinated manners. While many extramodal sensory cues compete for cognitive resources in these ecological contexts, that rodents can engage in such odor-guided behaviors suggests that they selectively attend to odors. We developed a behavioral paradigm to reveal that rats are indeed capable of selectively attending to odors in the presence of competing extramodal stimuli and found that this selective attention facilitates accurate odor-guided decisions. Further, we uncovered that attention to odors adaptively sharpens their representation among neurons in a brain region considered integral for odor-driven behaviors. Thus, selective attention contributes to olfaction by enhancing the coding of odors in a manner analogous to that observed among other sensory systems.

## Introduction

From neonatal attachment and suckling responses (*1, 2*), to selecting mates, finding food sources, and avoiding predators (*3–5*), rodent behavior is guided by odors in remarkably well-coordinated manners. The fact that these behaviors can be successfully orchestrated lends reason to believe that rodents must selectively attend to odors in these contexts at the expense of competing extramodal cues. For instance, as a rat forages for food, it must simultaneously ‘filter’ out competing auditory and visual stimuli arising from irrelevant sources. Rodents readily display shifting of attentional sets, including those involving odors (*6*), and can display attention towards information from other modalities (*7, 8*). It is entirely unknown, however, if selective attention regulates olfactory perception in rodents. This question is of great importance given the prevalence of rodents as models for olfactory function and due to the powerful control of olfactory perception by attention in humans (*9, 10*).

Given the aforementioned relevance of olfaction for survival, we reasoned that the olfactory system adaptively encodes odor information in manners dependent upon attentional demands. This would provide a mechanism for the brain to adjust the processing of incoming odor information based upon behavioral demands and context. We predicted that selective attention would shape the representation of odors within the ventral striatum (VS). This is likely given that the VS is important for evaluating sensory information in the context of motivated behaviors (*11*), a function considered integral for attention (*12*). Offering precedence for this is evidence provided by human functional imaging for increased hemodynamic responses to odors in the VS during attention (*9, 13*), particularly in the olfactory tubercle region of the VS, which is extensively innervated by olfactory input (*14*). The coding strategy VS neurons engage in, which may underlie this phenomenon, is unknown. Notably, the olfactory system does not have a classic thalamic relay, a component widely considered to be integral to attention and sensory awareness in other systems (*15*). The VS does, however, receive input from a variety of frontal cortex and neuromodulatory systems (*11*), which may allow for attention to sculpt odor coding. Defining the control of olfactory processing by selective attention has been hindered by a complete lack of a behavioral task to precisely manipulate odor-directed selective attention in rodents. Here we developed such a task, and combined it with single-unit neural recordings, to uncover fundamental principles of how rats utilize selective attention in manners advantageous for olfactory decision making. Our results indicate that selective attention to odors facilitates engagement in accurate olfactory decisions and enhances the contrast of odor representation in the VS by amplifying odor signal-to-noise ratios.

## Results

We first sought to demonstrate that rats are capable of displaying selective attention to odors. We developed a behavioral paradigm termed the two-alternative forced choice olfactory attention task (2AC-OAT; **Fig 1A&B**). The 2AC-OAT is a modified version of a standard 2AC task, wherein rats nose-poke into a center port and receive a stimulus that signals reward availability in either of two neighboring side ports. In the 2AC-OAT, across pseudorandom trials, rats were shaped to discriminate between two odors, and separately, to detect the absence or presence of a tone. Then, both tones and odors were presented simultaneously, and the rats learned to selectively attend to only one of the modalities to retrieve rewards.

**Fig. 1.**
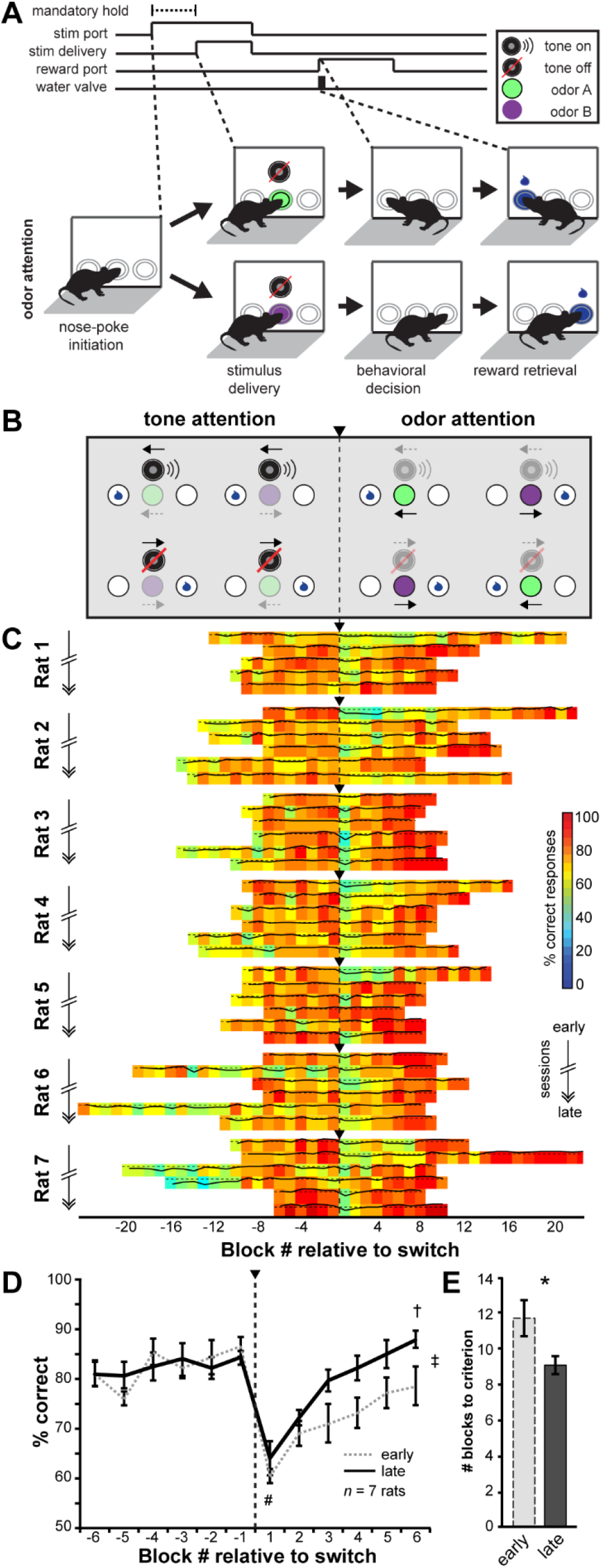
Odor-directed attention dictates discrimination accuracy. **(A)** 2AC-OAT trial outline. Example trials show correct 2AC choices during odor-directed attention on ‘tone off’ trials. Dashed line indicates mandatory preparatory hold time. **(B)** Four possible trials during the final phase of the 2AC-OAT. Top arrows for each trial indicate reward direction for tone cues; bottom arrows indicate reward direction for odors. Faded icons indicate cues that are present, but should be ignored when attending to the correct modality. **(C)** Example 2D histograms displaying performance of 7 rats over the course of six sessions of switching their attention during the 2AC-OAT (20 trials/bin). Solid overlaid lines indicate performance (% correct); dashed horizontal lines indicate criterion performance (80%). Vertical dashed line with arrowheads indicates the uncued experimenter-controlled switch from tone to odor attention. See Materials and Methods for additional details. For each rat, the top three rows were from early sessions, the bottom three rows from late sessions, except rat 1 which has 5 sessions. **(D)** Average performance of all 7 rats relative to the attentional shift, on their first two (early) and last two sessions (late). Performance dropped to chance levels immediately after the task switch and returned to criterion as the rat shifted its attention to odors. Note that performance improved more quickly during late sessions. **(E)** The average number of blocks for each rat to reach criterion after the attentional shift (6 blocks ≥80%) for early and late sessions. ^#^*p*<0.0001 (block -1 vs. 1, late), ^†^*p*<0.01 (block 1 vs. 6, late), ^‡^*p*<0.05 (block 6 early vs. block 6 late), ^*^*p*<0.05; two-tailed, paired t-test.

Several important features distinguish the 2AC-OAT. First, it provides robust, controlled stimulus presentations by requiring animals to nose poke to await stimuli. Second, it is an operant task, wherein several hundreds of trials can be completed within a single session, throughout which all conditioned stimuli are assigned equal valence, with both modalities predicting reward availability at some time during the session. These features are not inherent in main-stream attentional set-shifting tasks, preventing their use to study olfactory selective attention (*e.g.,* (6)). Third, out of the four possible 2AC-OAT trial types, odors may be either unattended or attended (**Fig 1B**, ‘tone attention’ vs ‘odor attention’). Further, half of these trials do not include a tone (**Fig 1B**, bottom half of trials), which allows for direct comparisons of the same odor across a session while it is attended versus unattended, without multisensory confounds (unlike the Wisconsin-Card Sorting Task (*16*) or (*8*)). Fourth, rats perform both the single-modality 2AC odor discrimination and the more challenging multi-modal 2AC-OAT in the same session, which allows for questions related to task demand to be addressed. Finally, the attentional switch is not cued nor overtly anticipated by the rats, which eliminates cue-generated expectation and allows for behavioral flexibility and odor coding relative to the attentional switch to both be probed.

### Rats selectively attend to odors and this dictates discrimination accuracy

We shaped 7 water-motivated rats to perform the 2AC-OAT. Over several phases, rats were shaped to criterion performance (≥85% correct responses) on the 2AC odor discrimination and tone detection tasks, separately (**Figs S1A-E, S2**). We then introduced modality switching, alternating performance between the single-modality 2AC odor discrimination and tone detection tasks within a session (**Fig S1F**). Next, rats began the session performing the tone detection task, and after reaching criterion performance, we switched them to a variant of the odor discrimination task (**Fig S1G**). Here we continued to present the same tones simultaneously with the odors. The rats were faced for the first time with the task of selectively attending to odors in the presence of now-irrelevant tones. The simultaneously presented tone and odor cues were either congruent (non-competing, signaling the same reward-port side) or incongruent (competing, signaling the opposing reward-port side). Once criterion was achieved, the rats advanced to the final 2AC-OAT (**Fig 1B, S1H**). The first half of the session consisted of auditory attention blocks (‘tone attention’), wherein rats attended to the presence or absence of a tone, while the odors were presented simultaneously. After reaching criterion performance (≥80% correct responses/block for ≥ 6 blocks), the task was switched to ‘odor attention’ blocks, wherein rats now had to direct their attention to the conditioned odors, ignoring the competing auditory information to which they had previously been attending. This switch was not cued and the rats had to rely upon their behavioral response feedback (reward receipt or not) to assess that the task contingencies had switched. It took the rats 392.6±44.6 blocks, across 24.9±1.3 sessions, to reach the first criterion switch for this final phase of the 2AC-OAT (**Tables S1-4**). Numerous successive sessions of over-training were given to establish robust behavioral performance. Among the last four sessions of this over-training, the rats took an average of 10.5±0.8 and 9.7±0.5 blocks to reach criterion for the tone attention and odor attention tasks, respectively.

Several significant findings emerged from the rats’ 2AC-OAT performance. First, we found that task accuracy is dependent upon the animal’s attentional strategy. Following shaping, rats performed the ‘tone attention’ task, despite the presence of competing conditioned odors, with an average of 85.48% correct responses per block (±1.14 SEM, inter-animal range: 82.92-91.25%). Directly after the task was changed from ‘tone attention’ to ‘odor attention,’ there was an immediate decrease in performance (*t*(6)=9.78, *p*<0.0001, block -1 vs block 1; **Fig 1C&D**). The rats initially made perseverative errors, reflective that they maintained their strategy of attending to the tone. As they received feedback on their errors (no reward), the rats modified their strategy and began directing their attention to odors, which consequently led to increased task accuracy (*t*(6)=-4.88, *p*=0.0028, block 1 vs block 6; **Fig 1D,** bold line). Rats displayed an average of 89.7% correct responses for ‘odor attention’ (±1.63 SEM, inter-animal range: 85.42-96.67%). Second, we observed that odor-directed selective attention is subject to plasticity with experience. Across sessions, rats improved their ability to shift their attention to odors (**Fig 1C&D**, compare dashed vs. bold lines), with high levels of performance reached sooner in late sessions versus early sessions (*t*(6)=-2.74, *p*=0.034, block 6 (early) vs block 6 (late); **Fig 1D**). Rats took fewer blocks to reach criterion in late sessions (*t*(6)=3.34, *p*=0.016; **Fig 1E**), demonstrating that they switched their attention to odors more rapidly with experience. We also tested a subset of rats for their abilities to direct selective attention to odors when perceptual demands were increased, given that there is interplay between attention, performance accuracy, and perceptual difficulty in other sensory systems (*e.g.,* (*17*)). As odor intensity was reduced, rats required more blocks to shift their attention to odors (**Fig S3**). Further, in agreement with the known influence of attention in dictating subtle, yet critical aspects of behavior (*18, 19*), we also uncovered that trial congruency and multisensory input impact 2AC-OAT performance (**Fig S4**). Together, these results demonstrate that rats can selectively attend to odors and that odor-directed attention improves with experience.

### Attention controls the neural representation of odors

The finding that selective attention is a prerequisite for accurate olfactory decisions indicates that attention may also sculpt the neural representation of an odor in a manner that facilitates perception. Does the brain represent an odor, of equal intensity and valence, differently dependent upon whether it is attended? To address this question, rats were unilaterally implanted with drivable tetrodes (*20*) into the olfactory tubercle (OT) region of their VS. Not all implanted rats contributed physiology data due to electrode placement errors, poor signals, or their inability to perform the cognitively demanding 2AC-OAT following surgery. We successfully performed OT single-unit recordings from 4 rats (**Fig S5**), which also contributed data during 2AC-OAT performance (**Fig 1**). Across multiple recording sessions/rat (range: 6-10), we lowered the tetrodes, and identified 232 cell-odor pairs (116 total single-units x 2 odors) (**Table S5**).

To directly test for attentional modulation of odor coding, controlling for possible multisensory influences, we only analyzed ‘tone off’ trials (50% of trials) (**Fig S1H**, blue box). We identified four epochs relative to stimulus onset, to assess behaviorally-relevant changes in neuron firing: (1) background (-1400 to -800ms), (2) stimulus port approach (-800 to -600ms), (3) preparatory hold (-600 to 0ms), and (4) odor stimulus duration (0 to 400ms). Of particular interest is the odor epoch, during which information may be mapped onto the neurons and used to guide behavioral choice. Across the entire population, during odor attention, 36 neurons were modulated by odor (32.03% of 116). We identified 55 odor-modulated cell-odor pairs out of the 232 possibilities (23.71%, 116 x 2 odors); *n*=27 odor-excited, *n*=28 odor-inhibited, *n*=177 unmodulated, with some cells modulated by both odors (see Materials and Methods).

We found that odor-directed selective attention bi-directionally sculpts the coding of odors in the OT by increasing the FRs of odor-excited cell-odor pairs (**Fig 2A, S6A**), while further decreasing the FRs of odor-inhibited cell-odor pairs (**Fig 2B, S6B**) during the preparatory hold and odor epochs. To statistically represent significant FR changes for the cell-odor pairs, we classified the data using auROC analyses (*21, 22*) (see Materials and Methods), which represents changes in FR within sliding windows of time relative to a shuffled background distribution. Greater significance emerges during odor attention for both populations during the hold and odor epochs (**Fig 2C&D**). During odor attention, for odor-excited cell-odor pairs, a large proportion of the population was significantly and rapidly excited during the hold and odor epochs (**Fig 2E**). The duration of this excitement was significantly longer during both the preparatory hold and odor epochs when rats attended to odors versus when they attended to tones (hold: *t*(26)=-3.20, *p*=0.0036; odor: *t*(26)=-3.51, *p*=0.0016), while 2AC odor discrimination was not significantly different (hold: *t*(26)=-2.15, *p*=0.041; odor: *t*(26)=-2.37, *p*=0.026) (Bonferonni critical *p*=0.0167; **Fig 2F**). Similarly, for odor-inhibited cell-odor pairs, a large proportion of the population was significantly and rapidly inhibited during the hold and odor epochs (**Fig 2G**). The duration that odor-inhibited cell-odor pairs were significantly suppressed relative to background during both the preparatory hold and odor epochs was significantly increased during odor attention as compared to tone attention (hold: *t*(27)=-3.79, *p*<0.001; odor: *t*(27)=-4.09, *p*<0.001) and odor discrimination (hold: *t*(27)=-3.87, *p*<0.0001; odor: *t*(27)=-4.67, *p*<0.0001) (Bonferonni critical *p*=0.0167; **Fig 2H**).

**Fig. 2.**
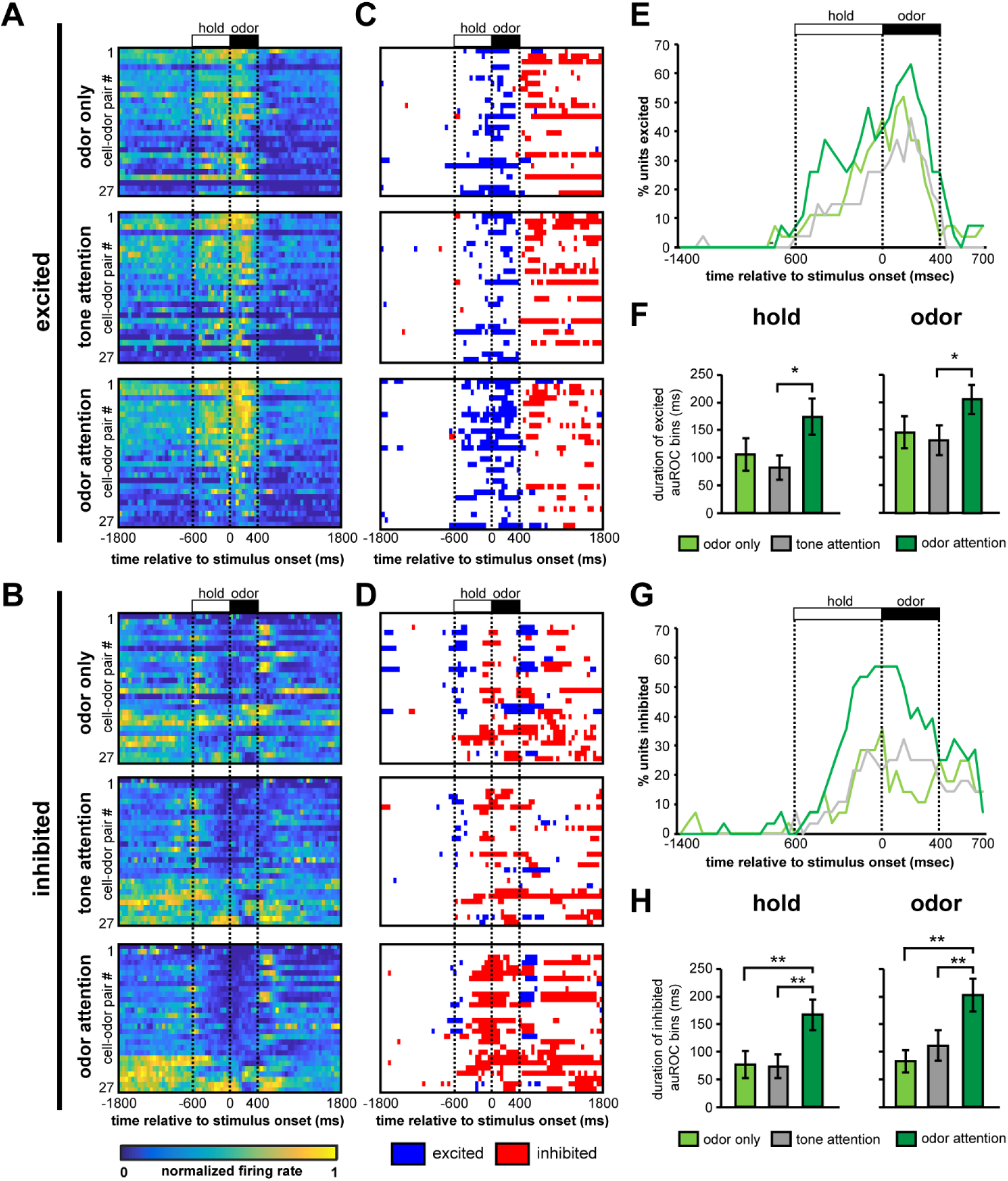
Odor-directed attention controls odor coding. 2D histograms (50ms bins) displaying normalized firing rates (FRs) of odor excited **(A)** and odor-inhibited **(B)** cell-odor pairs across the task states. Units are arranged from highest to lowest FRs, averaged over the first five bins post-stimulus onset during odor attention. See Materials and Methods for normalization details. As indicated by auROC significant bins, odor attention increases the FRs for odor-excited cell-odor pairs **(C)**, and further decreases the FRs for odor-inhibited cell-odor pairs **(D)** during the preparatory hold and odor epochs. Each row represents the corresponding neuron from the 2D histograms in **A** and **B.** Odor-directed attention increases the percentage of odor-excited cell-odor pairs that have significantly excited activity (**E**) and the percentage of odor-inhibited cell-odor pairs that have significantly inhibited activity (**G**) relative to background, earlier and for a longer duration. Odor attention thus significantly increases the duration of excitement (**F**) and inhibition (**H**) during both hold and odor epochs. ^*^*p*<0.05, ^**^*p*<0.01, two-tailed, paired *t*-test. Data from four rats (same as in **Fig 1**), 2-6 sessions/rat.

The above results indicate that selective attention to odors bi-directionally controls both OT ensemble activity and the representation of odors. To define how individual neurons incorporate attentional demands into their representation of odors, we used the cell-odor pairs classified above and calculated their change in FR with attention (ΔHz_attention_=FR_attended_-FR_unattended_) to yield a simple index for the direction of change in firing. Neurons were classified, for each epoch, as shifted negatively or positively if their FR either increased or decreased ≥1Hz. Among those odor-excited cell-odor pairs whose FRs shifted, we found that the majority decreased their background FRs (70%, 7/10), while increasing their FRs during the hold (70.6%, 12/17) and odor (60.0%, 12/20) epochs with odor-directed attention (**Fig 3A**). The proportion of odor-excited cell-odor pairs with decreased background FRs was greater than the proportion with increased background FRs (One sample proportion z=2.8, *p*<0.01), while the proportion of cell-odor pairs with increased FRs during the preparatory hold was greater than the proportion with decreased firings rates (z=3.7, *p*<0.001). The proportion of cell-odor pairs with increased FRs during odor did not reach significance (z=1.8, p=0.0679).

**Fig. 3.**
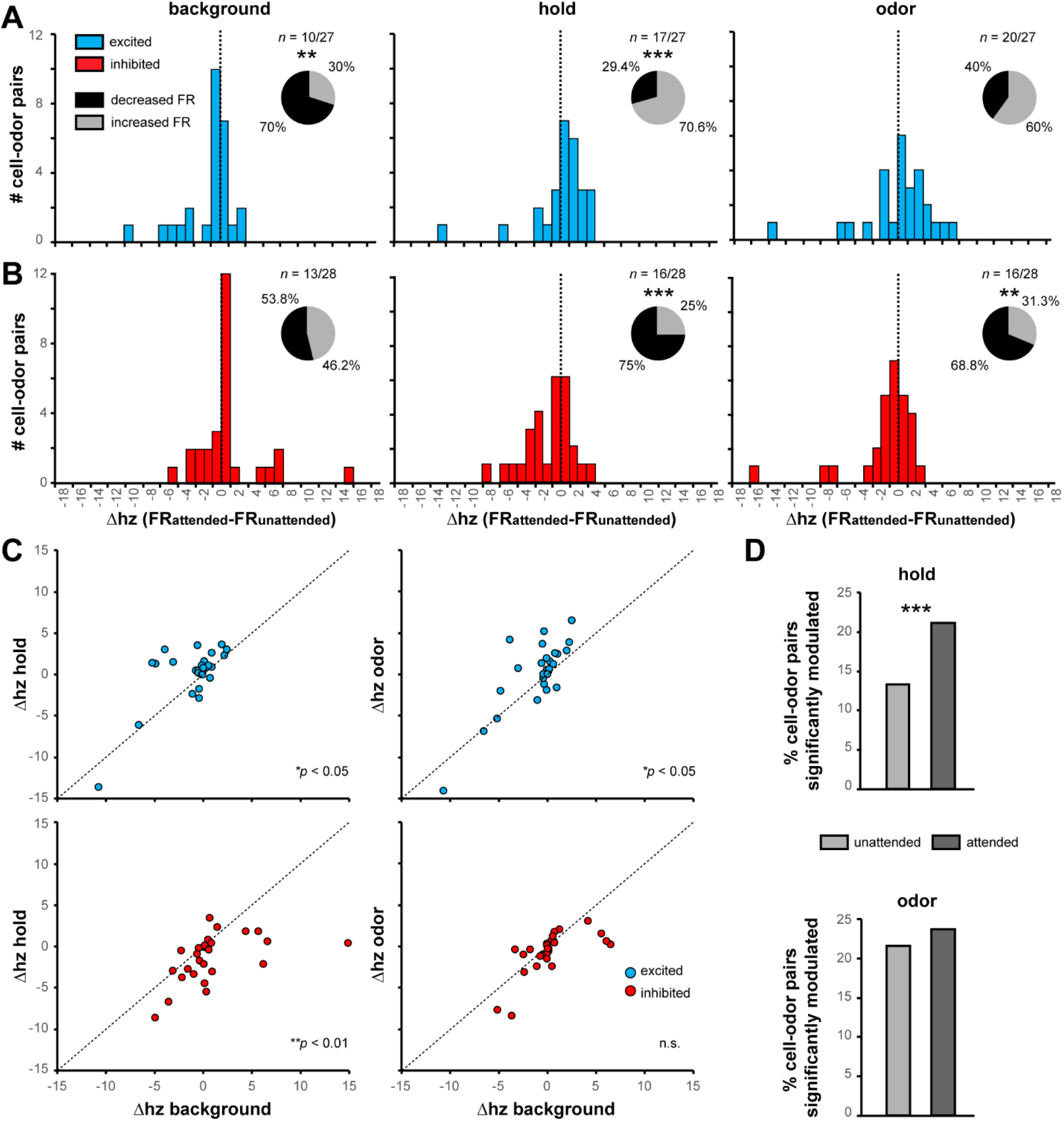
Attention yields enhanced signal-to-noise among odor coding neurons. Changes in FR with odor-directed attention, ΔHz_attention_=FR_attended_-FR_unattended_, for odor-excited **(A)** and odor-inhibited **(B)** cell-odor pairs for the three behavioral epochs. Pie chart: The proportion of increased or decreased FRs among units that shifted either negatively or positively (≥1Hz). **(C)** ΔHz_attention_ during background plotted against ΔHz_attention_ during either the hold (left) or odor (right) epochs for excited (top) or inhibited (bottom) unit populations. Excited cell-odor pairs falling above the dotted line indicate a greater change in FR relative to background. Inhibited cell-odor pairs falling below the dotted line indicate a greater decrease in FR relative to background. **(D)** The percentage of cell-odor pairs classified as either significantly excited or inhibited relative to background during the specified epochs are increased with attention to odor, particularly during the preparatory hold. Data from rats and sessions as in **Fig 2.** *ns* above pie charts indicate the number of cell-odor pairs shifted out of the total number of excited or inhibited cell-odor pairs. *p*<-values in **(C)**, as denoted, are two-tailed, paired *t*-test. ^**^*p* <0.01, ^***^*p* <0.001, one-proportion z-test.

An opposite direction of change was observed among the odor-inhibited cell-odor pairs, where among those whose FR changed, the majority decreased their firing during the hold (75.0%, 12/16) and odor (68.8%, 11/16) epochs while the rats were attending to odors (**Fig 3B**). A greater proportion of odor-inhibited cell-odor pairs decreased their FRs during the preparatory hold (z=4.6, *p*<0.0001) and odor epochs (z=3.2, *p*=0.0012) with attention. Notably, we determined that these effects were selective to odor-modulated cell-odor pairs, as the majority of FRs for those which were unmodulated were unchanged during background (87.01%, 154/177), hold (88.70%, 157/177), and odor epochs (87.57%, 155/177; **Fig S7A**). Among those unmodulated cell-odor pairs that were shifted (11.30%, 20/177), we did observe that a greater proportion displayed decreased firing during the preparatory hold (z=3.9, *p*< 0.0001). Overall, with odor-directed attention, odor-excited cell-odor pairs display enhanced firing relative to background in preparation for the upcoming stimulus, while odor-inhibited cell-odor pairs show decreased firing relative to background in preparation for and during the odor stimulus.

We reasoned that odor-directed attention may control individual neural FRs in two ways. For a given neuron, the overall FRs throughout all epochs may be broadly influenced in direction and magnitude by odor attention, indicative of a general ramping up or down of overall activity. Alternatively, as suggested by **Figures 3A and B**, odor attention may control odor signal-to-noise ratios. Along this logic, for an odor-excited neuron, activity within the preparatory hold and odor epochs may be increased, while background activity remains either unchanged or is decreased. For an odor-inhibited neuron, activity during the preparatory hold and odor epochs may be further suppressed, while background activity remains either unchanged or is increased.

To address the above question, in a final series of analyses, we compared the ΔHzattention of the background to either the hold or odor epochs for both odor-excited and odor-inhibited cell-odor pairs (**Fig 3C**). Points falling along the unity line would indicate FR changes that are similar in direction and magnitude across the epochs, which would support a general change in neural activity within a trial, irrespective of epoch-specific influences. We found, however, for odor-excited cell-odor pairs, that the change in FR during both the preparatory hold and odor epochs was increased relative to the change in FR of the background (hold: *t*(26)=-2.32, *p*=0.028, odor: *t*(26)=-2.54, *p*=0.017) **Fig 3C**, top). In many cases, the background FR decreased, while the FR during the hold and odor epochs increased. Furthermore, for odor-inhibited cell-odor pairs, the change in FR during the preparatory hold period was more greatly decreased relative to the change in background FR (hold: *t*(27)=3.227, *p*=0.003, odor: *t*(27)=1.93, *p*=0.064; **Fig 3C**, bottom), and thus we conclude that odor-directed attention enhances the signal-to-noise within these cell-odor pairs. Notably, this effect is specific to odor-modulated neurons, since those classified as unmodulated by odors during odor attention displayed FR changes that were similar in both their direction and magnitudes (hold: *t*(176)=1.38, *p*=0.169, odor: *t*(176)=1.01, *p*=0.313; **Fig S7B**). Consequently, odor-directed attention recruited more cell-odor pairs to encode the acts of the preparatory hold (odor attention: 21.12%, 49/232 vs tone attention: 13.36%, 31/232) and odor sampling (odor attention: 21.552%, 50/232 vs tone attention: 23.707%, 55/232) (**Fig 3D**). Therefore, our results indicate that selective attention facilitates odor coding within task-critical moments by enhancing the signal-to-noise ratio of odors.

## Discussion

We have demonstrated that rats are capable of selectively attending to odors in the presence of conflicting stimuli. We predict this executive capacity affords rodents the ability to engage in ecologically critical behaviors (*e.g*., foraging, predator avoidance, mate selection), which are highly multisensory contexts requiring animals to focus at times upon a single modality at the expense of others. Not only does our work show that selective attention enhances odor discrimination capacity, but also that this ability improves with experience. This result highlights an important interplay between attention, olfactory processing, and learning and indicates that rodents develop a strategy for selectively attending to odors.

Equally important is our finding that selective attention contributes to olfactory processing by enhancing the contrast of odor representation in the OT by amplifying odor signal-to-noise ratios. This sculpting of odor information by attention is analogous to that observed upon attentional modulation in the visual and auditory systems of more cognitively-advanced mammals (*e.g*., (*17, 23, 24*)). We predict that this function, together with possible attentional modulation in other olfactory structures, is likely responsible for the effect of selective attention on facilitating accurate odor discrimination. The OT is among several structures that together make up the olfactory cortex. Therefore, attention may ‘filter’ available odor information into the entirety of down-stream structures important for emotion, motivation, and memory. Moreover, this discovery lends credence to an important question: what system is responsible for this attentional state-dependent control of olfaction given that there is no mandatory thalamic relay in the olfactory system? Indeed, the OT neurons we recorded from herein receive the bulk of their input directly from the brain’s initial odor processing stage, the olfactory bulb, and also from the neighboring piriform cortex (*14, 25*). While the olfactory bulb is hypothesized to serve functions analogous to the thalamus (*26*), attentional modulation may also arise from frontal cortex afferents (*27*) or from descending neuromodulatory influences upon these neurons (*11, 28*).

Taken together, a rodent, just like a human (*9, 10*), can employ selective attention to aid in olfactory perceptual goals and this attention enhances the representation of odor information within part of a brain system that is integral for evaluating sensory information in the context of changing motivational demands. Our results put forward a model whereby an attention-dependent signal-to-noise coding strategy facilitates odor perception.

## Acknowledgements

This work was supported by NIH NIDCD grants R01DC014443 and R01DC016519 to D.W. and F31DC014615 to K.C. We thank Dr. Ben Strowbridge for helpful discussions throughout this study.

